# Attention Decorrelates Sensory and Motor Signals in the Mouse Visual Cortex

**DOI:** 10.1101/615229

**Authors:** Mohammad Abdolrahmani, Dmitry R Lyamzin, Ryo Aoki, Andrea Benucci

**Affiliations:** RIKEN Center for Brain Science, Wako-shi, 351-0198 Saitama, Japan; University of Tokyo, Graduate School of Information Science and Technology, Department of Mathematical Informatics, 1-1-1 Yayoi, Bunkyo City, Tokyo 113-0032, Japan

**Keywords:** Vision, decision-making, mouse, behavior, movement-related activity

## Abstract

Visually-guided behaviors depend on the activity of cortical networks receiving visual inputs and transforming these signals to guide appropriate actions. However, non-retinal inputs, carrying motor signals as well as cognitive and attentional modulatory signals, also activate these cortical regions. How these networks avoid interference between coincident signals ensuring reliable visual behaviors is poorly understood. Here, we observed neural responses in the dorsal-parietal cortex of mice during a visual discrimination task driven by visual stimuli and movements. We found that visual and motor signals interacted according to two canonical mechanisms: divisive normalization and response demixing. Interactions were contextually modulated by the animal’s state of attention, with attention amplifying visual and motor signals and decorrelating them in a low-dimensional space of neural activations. These findings reveal canonical computational principles operating in dorsal-parietal networks that enable separation of incoming signals for reliable visually-guided behaviors during interactions with the environment.

## Introduction

Dorsal-parietal areas are indispensable for visual perception and visually-guided behaviors, but only a small fraction of their synaptic inputs originate from the eyes. Even the primary visual cortex (V1) receives less than 5% of its inputs from the eye (Ahmed et al., 1994; Binzegger et al., 2004). Most inputs are local, from within the cortex, or long-range, from other brain regions (Crapse and Sommer, 2008; Ibrahim et al., 2016; McCormick et al., 2015; Niell and Stryker, 2010; Sommer and Wurtz, 2008; Stringer et al., 2019; Zhang et al., 2014). Traditionally, the influence of extra-retinal inputs on visual processing has been minimized with the aid of anaesthetized, paralyzed preparations. However, due to advances in recording techniques in awake behaving animals, a growing number of studies have demonstrated that extra-retinal signals play a fundamental role in the contextual modulation of visual signals during goal-directed behaviors. For instance, recent studies conducted in rodents have shown that locomotion enhances visual responses (Fu et al., 2014; Niell and Stryker, 2010), modulates spatial integration (Ayaz et al., 2013), improves neural encoding (Dadarlat and Stryker, 2017), and affects the processing of visual information in a task-specific manner (McBride et al., 2019). Besides locomotion activity, other afferents to dorsal-parietal cortices contextually modulate visual responses. Variability in internal brain states associated with cognitive variables (e.g., attention, alertness, task engagement, and arousal) affect visual coding and perception (McGinley et al., 2015a) and are distinct from locomotion-related activity (Vinck et al., 2015). For instance, visual selective attention—also reported in mice (Wang and Krauzlis, 2018)—can modulate visual responses via long-range top-down connections from frontal cortical areas (Zhang et al., 2014).

Taken together, current physiological and computational observations suggest that dorsal-parietal networks are a multimodal processing system that integrates visual, motor, and cognitive-related signals (Saleem et al., 2013). However, a key unanswered question is how these signals concurrently impinging on the dorso-parietal networks interact with each other for reliable visual behaviors.

We examined this question by recording GCaMP responses across a large network of dorsal-parietal areas as mice engaged in a visual discrimination task. We found that movement signals are large in amplitude and broadly activate these cortical regions, consistent with previous reports (Parker et al., 2020), finding also that movement responses activated visual cortices regardless of the retinotopic area boundaries and disregarding principles of retinotopic activations. Using a GLM model and focusing on saccadic and body movements, we established that movements summed supra-linearly with visual responses, but sub-linearly with each other, consistent with the mechanism of divisive normalization.

Notably, these interactions were contextually modulated by the state of sustained attention of the animal, with visual and motor signals both amplified during heightened attention. Overall, these effects could be represented in a low-dimensional activity space of dorsal-parietal areas, with decorrelated stimulus and movement trajectories reflecting the non-retinotopic characteristics of movement signals and with the decorrelation of signals becoming stronger in heightened states of attention.

## Results

### Movement-related responses in the visual cortex

Mice (n = 10) were trained in a two-alternative forced choice (2AFC) orientation discrimination task (Figures 1A, 1B, S1A, Methods) with a high-throughput automated system featuring self-head fixation (Aoki et al., 2017). Rotations of a toy wheel controlled by the front paws of the animals produced horizontal shifts of two grating stimuli presented on a screen positioned in front of the animal. The task was to shift the most vertical of the two stimuli to the center of the screen to obtain a water reward. Besides wheel rotations, during the task mice made saccadic eye movements mostly in the nasal-temporal direction and preferentially after the stimulus presentation (open loop, OL, Figures 1C, S1B; Movie S1). As mice engaged in the task, we imaged responses from excitatory neurons in a large network of visual cortical areas (Figures 1D, S1C). Notably, responses to isolated saccades were about four times larger than responses to visual stimuli even in the V1 retinotopic location of the stimulus (Figure 1E, 2.5 ± 0.4% dF/F peak amplitude for saccades vs 0.57 ± 0.05% dF/F for stimulus, p<10^−4^, t-test), and in agreement with recent reports (Musall et al., 2019). Response amplitudes increased proportionally to saccade size (Figure 1G). Saccades strongly activated medial V1 and the medial part of the dorsal visual stream (areas PM, AM, A)—regions implicated in the stabilization of visual perception during ocular movements (Ross et al., 1996; Wurtz et al., 2011)—as well as anterior visual-parietal areas (Lyamzin and Benucci, 2019).

**Figure 1.**
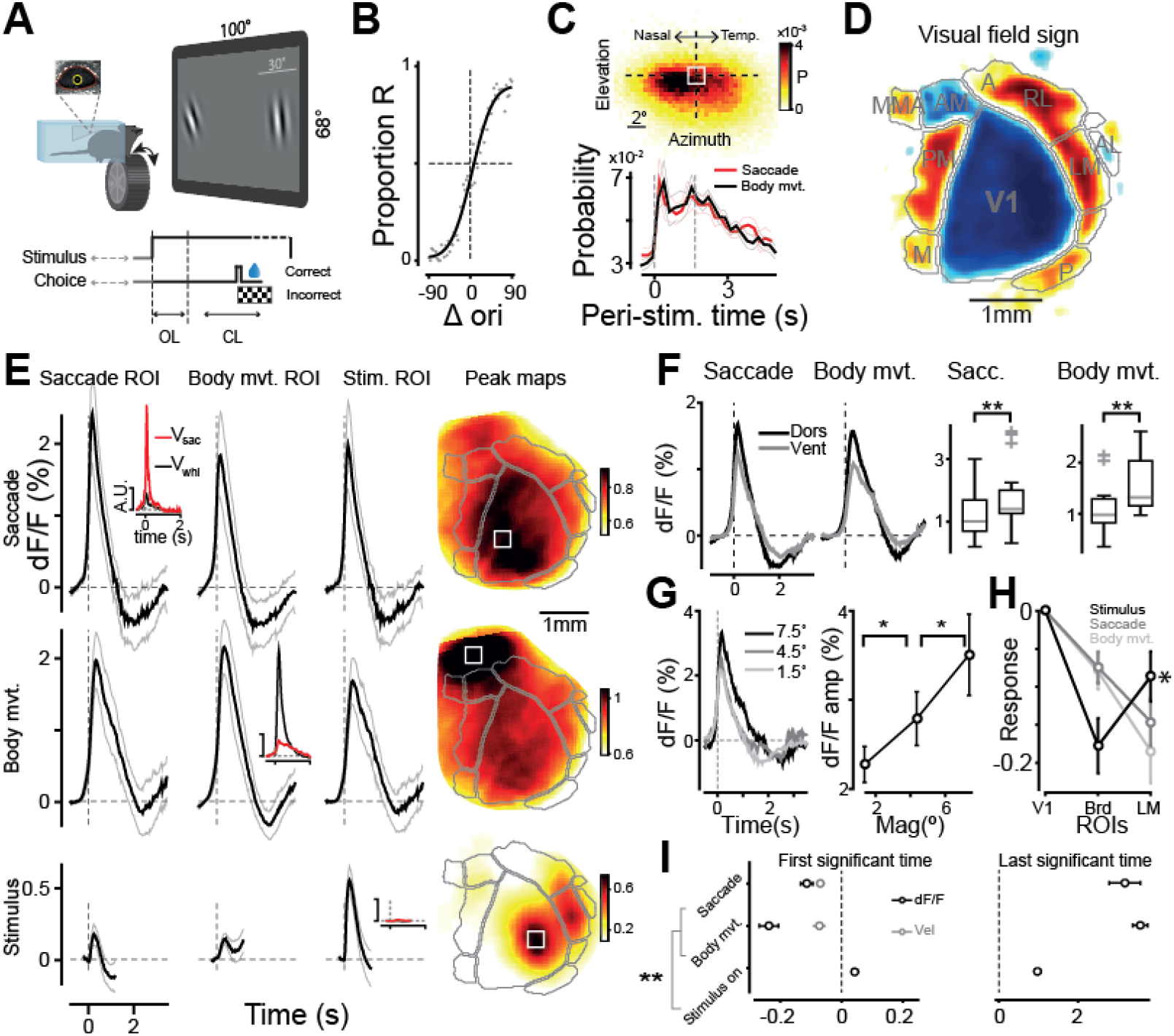
Neuronal responses to saccadic and body movement are non-retinotopic. (A) Mice are trained in a 2AFC orientation discrimination task. Animals reported their choice by rotating a wheel with their front paws, and eye movements were simultaneously recorded (inset, Methods). (B) Mice learned the task after a few months (66 ± 14% correct; mean ± s.d., Methods). Psychometric curve for an example mouse: black line, fit; gray dots, data. (C) Eye movements during the task were mostly along the horizontal direction. Top: probability (P) of saccadic landing positions. White square, eye neutral position (Methods). Bottom: probability of detecting a saccade or a body movement during the trial is larger than before the trial (p < 10^−4^ Wilcoxon test) for both movement types; n = 10; thin lines s.e.). Probabilities for naïve mice are shown in Figure S1B. (D) Visual area borders are identified using field-sign maps (Methods, Figure S1C). (E) Responses to saccadic and body movements are about 4x larger than visual responses. Rows for event types and corresponding ROIs on the columns (ROIs are shown as white squares in the ‘peak maps’). Max amplitude of saccadic response: 2.5 ± 0.4 % dF/F ± s.e.; t-test comparing to peak stimulus amplitude, p < 10^−4^. FWHM 0.56 ± 0.06 s. Body-movement response: peak 2.2 ± 0.2; p < 10^−4^, t-test; FWHM 0.83 ± 0.07 s. Peak stimulus response: 0.57 ± 0.05%. Insets: velocities of eye movements (red) and body movements (black). Rightmost column: response maps at times of peak amplitude. (F) Left: Isolated saccadic and body movement responses averaged across dorsal (M, PM, AM, RL) and ventral (LM, P, POR) stream areas. Right: Amplitude difference between ventral and dorsal stream areas at the time of peak response for saccades and body movements: p = 0.002, p = 0.004 (t-test, n = 10). (G) Left: Saccadic responses to three different saccade amplitudes (1.5°, 4.5°, and 7.5°; shades of gray) in saccade ROI. Right: amplitude of saccadic responses at the time of peak response for different saccade magnitudes (p = 0.01, t-test, n = 10); error bars, s.e. (H) Stimulus response at the V1-LM border (Brd) is significantly smaller than in LM (p = 0.019, Wilcoxon test), but significantly larger for saccade (p = 0.002), and n.s. larger for body movement responses (p = 0.105, t-test). (I) First (left) and last (right) time points are significantly different from baseline activity in the [−0.8 −0.3] s window before saccade and body movement detection and in the [−0.3 0] s window for stimulus onset. Saccades, −110 ± 20 ms; body movements −237 ± 31 ms; n = 10. Gray dots, detection time of 1st significant time point for saccadic and body movement velocities (Methods). Pre-movement neural responses precede that of the actual movement velocities.

Wheel rotations—hereafter ‘body movements’—included general movements of the trunk, tail, snout, and whiskers (Supp. Video 2). They typically occurred after the stimulus onset during the open-loop period (Figure 1C). Consistent with previous results on locomotion in rodents (Parker et al., 2020), body movements elicited large-amplitude responses (2.2 ± 0.2% dF/F peak amplitude), about three times larger than responses to visual stimuli (0.57 ± 0.05% dF/F, t-test p<10^−4^) and were localized in medial V1 and dorsal visual-stream areas, such as the posterior parietal cortex (A, anterior-RL, and AM) (Figures 1E, 1F, S1I).

Importantly, neuronal responses to saccadic and body movements were not significantly affected by reafferent signals (Crapse and Sommer, 2008), as confirmed by several observations: (1) Activity patterns emerged before movement onset (Figure 1I, saccade: −110 ± 20 ms; body movement: −237 ± 31 ms; s.e.; n = 10; Figure S1D), possibly reflecting a pre-motor, preparatory component propagating to these areas (Pinto et al., 2019). (2) For saccades, removing the visual stimuli did not significantly affect spatio-temporal response patterns, whereas activations induced by the screen edges were significantly different from those of saccadic movements (Figures S1E, S1F). Furthermore, simulating ocular movements by jittering the stimuli produced significantly different response patterns (simulated saccades, Figure S1G). (3) Signatures of reafference could be found using a singular-value decomposition analysis (SVD) of movement responses (Figure S1H), having small amplitudes and localized primarily in lateral areas involved in the processing of stimulus motion (Juavinett and Callaway, 2015) (Figures S1H, S1J).

Overall, saccadic and body movement responses prominently contributed to the trial-to-trial variability of the responses in the visual cortex, as they were consistently several folds larger than contrast responses and displayed distinct spatial and temporal activity patterns. Both types of movement signals concurrently activated several visual areas, but importantly, their response localization did not follow retinotopic principles characteristic of visually-evoked responses. According to retinotopy, large-amplitude V1 medial and anterior activations are evoked by stimuli located in the lower part of the visual field at large eccentricities. Stimuli in this part of the visual field also evoke strong responses in lateral areas (LM, AL. Figure S1F), with a characteristic ‘mirroring’ of activations at the vertical meridian (Sereno et al., 1994). Movement-related responses did not agree with this mapping, monotonically decreasing in amplitude across the vertical meridian (Figure 1H), making their response patterns in all generality inconsistent with activations elicited by visual stimuli.

### Nonlinear response summation and normalization

The concurrent representation of visual and non-visual signals across overlapping networks raises the question of what principles govern their interaction dynamics.

We modeled neural activity as a second-order Wiener series expansion (Meyer et al., 2017; Park and Pillow, 2011) (GLM, Methods), thus representing interactions of visual and non-visual signals as a sum of their individual responses (linear terms, Figures 2A, 2B), and pairwise nonlinear (2^nd^ order) terms quantified by interaction kernels (Figure 2C). Stimulus interactions with saccades and body movements were positive and peaked at 0.52 ± 0.14 s (saccade) and at 0.5 ± 0.01 s (body movement) after the stimulus onset, resulting in a supralinear response summation (Figure 2D). By contrast, saccade-body movement interaction kernels were largely negative, resulting in a sublinear summation of responses, especially when a saccade followed a body movement (0.58 ± 0.05 s relative lag) (Figures 2E, S2B). We confirmed that this result was not a consequence of GCaMP signal saturation (Figure S2A). The negative component in saccade-body movement interaction and the positive component in visuomotor interactions, at the time of their peak amplitude, were broadly distributed across visual areas, with the strongest nonlinearity localized similarly to the peak response amplitude of the total saccade-body movement response and stimulus-movement responses, respectively (Figures 2D, 2E).

**Figure 2.**
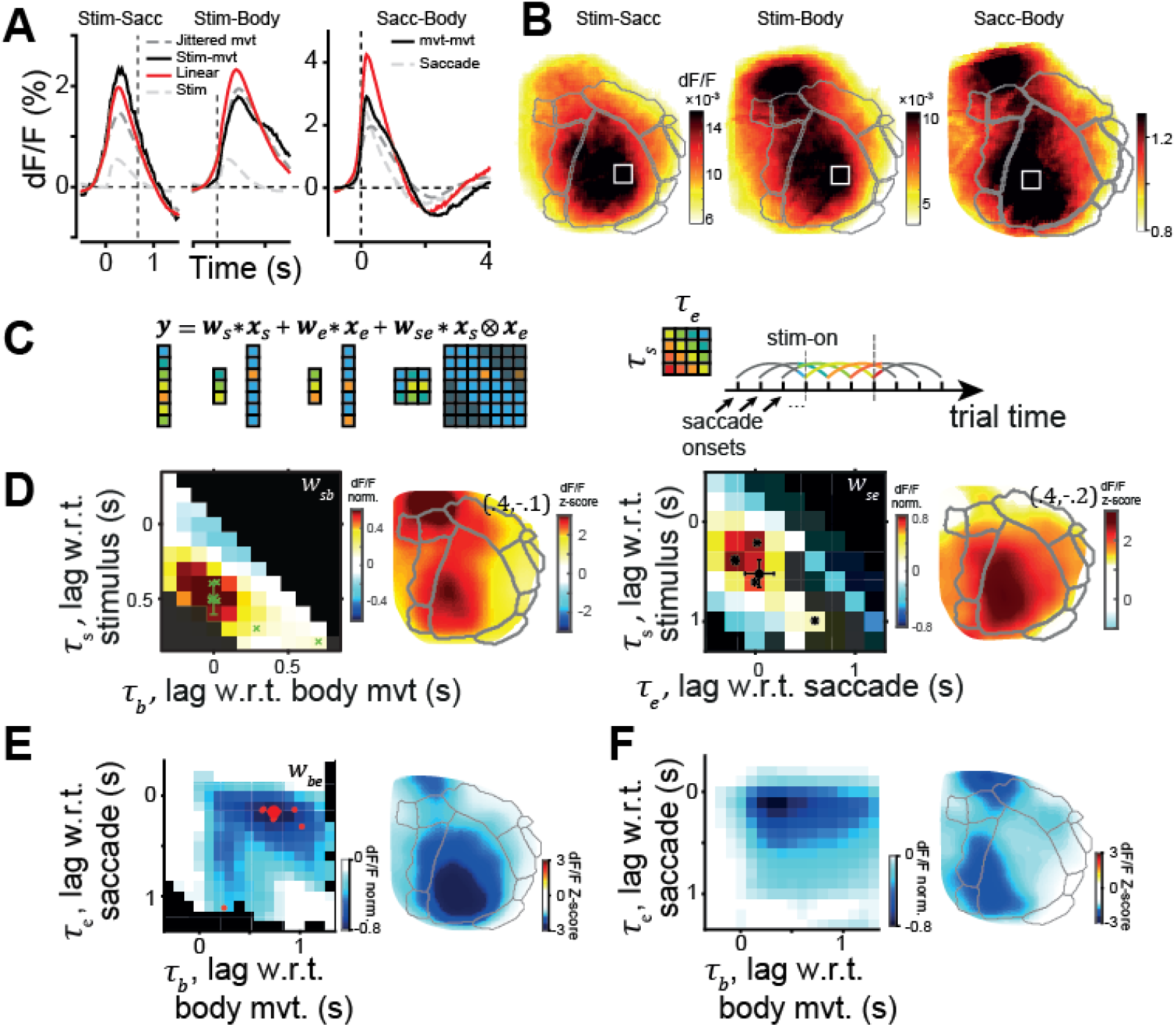
Nonlinear Response Summation and Normalization. (A) Linear prediction of stimulus-movement interactions: stimulus-saccade (left), stimulus-body movement interactions (middle), and saccade-body movement interactions (right) in the stimulus ROI (white squares in (B)). Linear model (red, Methods) and data (black). Gray lines are the responses to isolated stimulus (as in Figure 1E) and time-jittered saccadic or body movement responses (Methods). Peak response amplitude for stimulus-saccade (left) and stimulus-body movement interactions (middle), averaged across mice: 2.1 ± 0.22 dF/F (% ± s.e.) and 1.7 ± 0.14 dF/F, respectively. (B) Activity maps for panels in (A) at the time of peak response. (C) Left, GLM with an example stimulus-saccade nonlinear interaction term *W*_*se*_ (Methods). Briefly, inputs *x*_*s*_, *x*_*e*_ (*s*-stimulus, *e*-eye movement) are convolved (∗) with respective kernels *W*_*s*_, *W*_*e*_, and their outer product *x*_*s*_⊗*x*_*e*_ is convolved with *W*_*se*_ along the diagonal. Right: kernel values show a nonlinear part of pairwise interaction for two events of all relative lags (colors). Ticks: saccade onset times. Left dashed line: stimulus onset. (D) Left: normalized stimulus-body movement interaction kernel *W*_*sb*_, average of filtered and normalized kernels of (n = 6) mice. Black: unattainable kernel elements, dim-color elements are estimated from n<5 animals. Green crosses: maximum positive values of individual animals; large cross with error bars, their median (*τ*_*s*_ = 0.5 ± 0.1 s, *τ*_*b*_ = 0 ± 0.0 s, median ± median-derived s.d.). The activation map to the right shows the normalized model-predicted nonlinear component at lags (*τ*_*s*_, *τ*_*b*_) = (0.4, −0.1) s. Right. Normalized stimulus-eye movement interaction kernel *W*_*se*_, average of (n = 5) mice. Maximum positive values of individual animals are shown with black asterisks; population mean, black circle with error bars (*τ*_*e*_ = 0.04±0.15 s; *τ*_*s*_ = 0.52±0.14 s). The activation map to the right shows the normalized model-predicted nonlinear component at lags (*τ*_*s*_, *τ*_*b*_) = (0.4, −0.2) s. (E) GLM captures the sub-linearity of saccade-body movement interaction. Left: average saccade-body movement kernel (n = 8 mice, Methods). Masked black squares, contribution from less than five mice. Red dots: the largest negative values for individual animals; large circle for population average (lag of maximum nonlinearity relative to body movement *τ*_*b*_ = *t*_*maxNL*_ − *t*_*b*_ = 0.74 ± 0.06 s; relative to eye movement, *τ*_*e*_ = 0.16 ± 0.03 s; relative lag between the two movements, 0.58 ± 0.05 s); one outlier omitted, error bars smaller than circle size. Right: spatial distribution of the nonlinear component at the lag of the largest nonlinear interaction (large red circle in the left panel). (F) Same as (E), with GLM fitted to the output of the normalization model (Methods).

We examined the mechanisms underlying the sublinear summation of saccade and body movement responses with the aid of a divisive normalization model that could capture the dependence between the strength of the nonlinear interaction and the response amplitude (Carandini and Heeger, 2012). We implemented a model where the strength of the divisive scaling (normalization pool) was proportional to the product of the signals at each time point: the larger the signals, the stronger the divisive influence they exerted on each other (Methods). This one-parameter model largely explained the average sublinear summation, with the lag-dependent asymmetry explained by the differences in temporal response profiles between saccade and movement responses (Figure 2F).

Thus, interactions of stimulus and movement responses were supralinear at short temporal lags after stimulus onset, and interactions between saccadic and body movements were overall sub-linear, with a stronger sub-linearity when a saccade followed a body movement, according to a divisive normalization mechanism.

### Sustained attention modulates behavior and amplifies visuomotor signals

In trials with movements shortly after stimulus onset, pupil dilations were also the largest (Figures 3A, 3B), suggesting variability across trials in the internal state of the animal (McGinley, 2020). Modulations in internal states might depend, for instance, on the animal being more or less engaged or vigilant during the task. Hereafter, we refer to the broad spectrum of goal-directed internal states as to *sustained* attention (Havenith et al., 2018; Sarter et al., 2001; Unsworth et al., 2018). Across animals and behavioral sessions, larger pupil dilations following stimulus onset correlated with a tighter distribution of saccadic and body movement times (Figure 3A), possibly reflecting a learned hazard function (Luce, 1986) and attentiveness to the trial temporal structure. Pupil dilations were also predictive of trial outcome (Figure 3C). Furthermore, in a space defined by pupil area change and its baseline area, performance peaked at intermediate values (Figure 3C), suggesting hyperarousal states at very large pupil dilations in agreement with the Yerkes-Dodson inverted U-curve (McGinley et al., 2015b).

**Figure 3.**
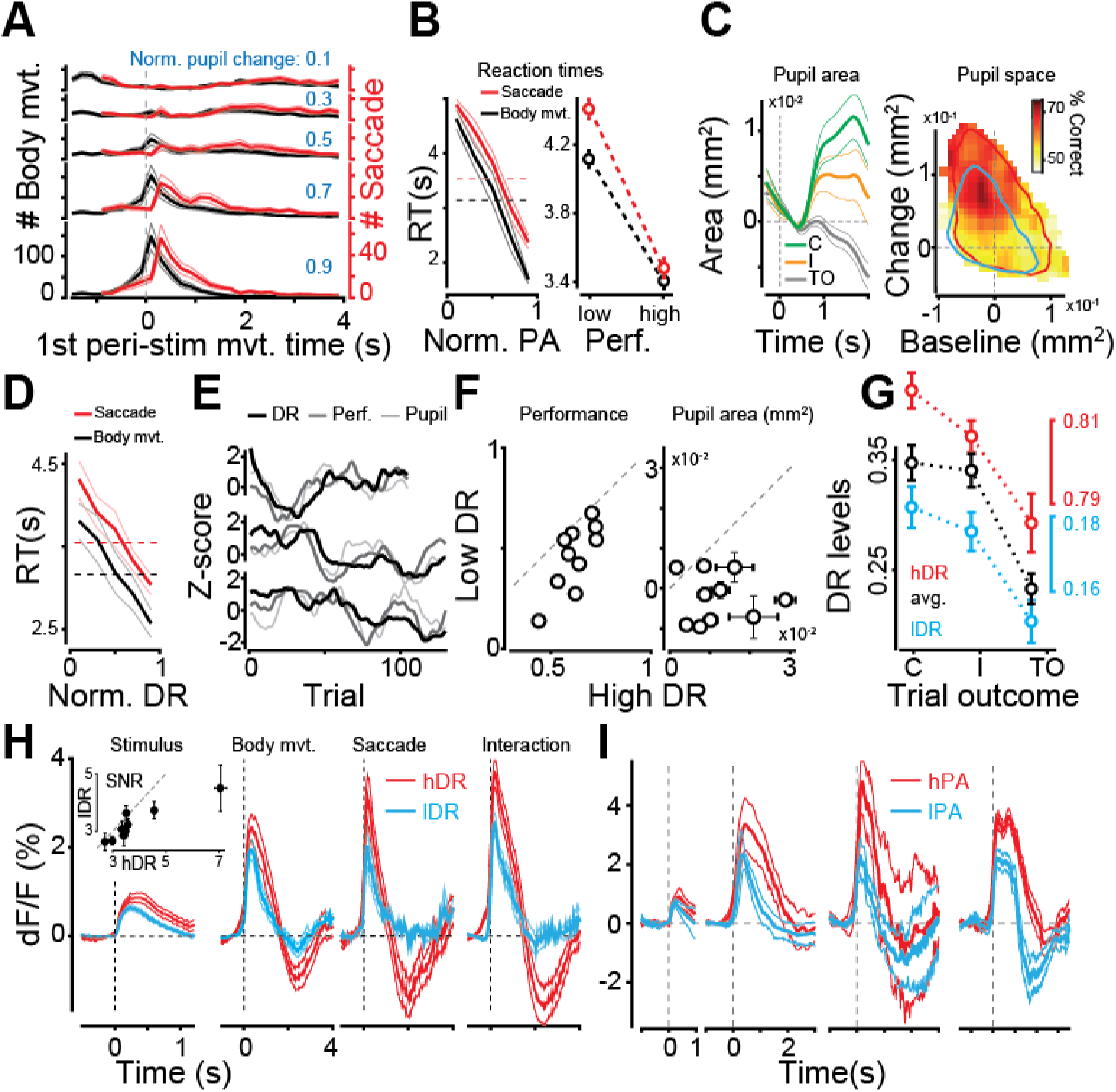
Sustained attention modulates behavior and amplifies visuomotor signals. (A) Histogram count of the first detected saccade (red) and body movement (black) relative to stimulus onset, in different attentional states, as indicated by the post-stimulus pupil-area change (blue text). (B) Trials with high performance and with a large pupil area are characterized by shorter reaction times (time of 1^st^ detected saccade or body movement after stimulus onset) (p < 10^−3^ saccade RT in high vs low performance trials, and in high vs low PA trials (p < 10^−3^, t-test for body movement RT in same comparisons). Horizontal dashed lines: average movement times across all pupil areas. (C) Pupil area correlates with trial outcome (traces aligned at t = 0.5 s). Left: pupil area changes after stimulus onset for correct (C), incorrect (I), and timeout (TO) trials. Peak area values in TO smaller than in C and I trials (p < 0.05, ANOVA), also in pairwise comparisons (C-TO, p = 0.0003; I-TO, p = 0.0461; n = 10, one-way ANOVA, corrected for multiple comparisons; C-I, p = 0.1132, n.s.). Right: performance in different regions of pupil space (Methods). Peak performance is at intermediate pupil dilations. Contour lines (90^th^ percentile) for trials with high (red) and low (cyan) DR index values. (D) Trials with a large DR index are characterized by shorter reaction times (time of 1^st^ detected movement after stimulus onset) (saccade p < 10^−4^; body movement p = 3·10^−4^, t-test). Horizontal dashed lines: average reaction times. (E) Performance fluctuations were correlated with changes in pupil area within blocks of several trials (example epoch, but n.s. when across all sessions: ρ = 0.02 ± 0.41, p = 0.37, t-test) and significantly correlated with DR index (ρ = 0.07 ± 0.03 p = 0.01, t-test, n = 270 session) (Methods, example sessions). (F) Trials with high DR index values are associated with higher performance (left) (p = 0.02, Wilcoxon test, n = 10) and larger pupil area (right) (p = 0.006, Wilcoxon test, n = 10). (G) Correct and incorrect trials have larger DR index values compared to timeout trials (C-TO, p = 10^−4^; I-TO, p < 10^−4^; n = 10, one-way ANOVA, corrected for multiple comparisons; C-I, p = 0.3213, n.s.). This tendency holds in different quantiles of the DR distribution: high DR (red, top 30%; C-I, p=0.0866; C-TO, p < 10^−4^; I-TO, p = 0.004) and low DR (blue, bottom 30%; C-I, p = 0.686; C-TO, p = 0.0003; I-TO, p = 0.0024). (H) Response amplitudes for isolated stimuli, saccadic and body movements, as well as during saccade-body movement interactions, are larger in states of high sustained attention (hDR) (all p < 10^−2^, Wilcoxon test). Inset: larger signal-to-noise ratio for isolated stimulus responses in high attention states (p = 0.02, Wilcoxon test). (I) same as (H), but trials are separated using pupil area. Peak response amplitudes of body movement, saccade, and saccade-body movement interactions were significantly larger in large pupil area trials (hPA) than in small pupil area trials (lPA) (p < 0.05, Wilcoxon test), and n.s. for stimulus response amplitude.

Sustained attention had a clear impact on cortical neural dynamics. We defined a measure of cortical state (dynamic range index, DR) that captured response variability (s.d. throughout trial time, Methods) across all cortical areas (Figures 3D–3I). This index significantly correlated with a previously introduced correlation-based measure of cortical state (Harris and Thiele, 2011) (Figure S3A). Changes in DR values correlated with modulations in reaction times, performance, pupil dilation, and trial outcome (Figures 3D–3G). During states of heightened attention (large DR values), the amplitude of the stimulus-evoked response and its signal-to-noise ratio increased (p<10^−2^, p = 0.02 respectively, Wilcoxon test) (Figure 3H, Methods) in agreement with other reports (Kozyrev et al., 2019; McAdams and Maunsell, 1999; Reynolds and Heeger, 2009; Treue and Martinez Trujillo, 1999). Both saccadic and body movement responses were also amplified during high DR states (Figure 3H) (p < 10^−2^, Wilcoxon test). A congruent result was found by using pupil dilations as a proxy for cortical state (p < 0.05, Wilcoxon test; n.s for visual stimulus), suggesting that response amplification did not follow from the definition of the index that measured response variability over time, but reliably indicated a *global* change in the state of the animal (Figure 3I) (Methods).

### Attention enhances the demixing of visuomotor signals

We next examined the differential impact of this amplification across cortical regions. We represented activations across all visual areas in an attention-invariant space of activity modes (Allen et al., 2019; Li et al., 2016). This defined a characteristic “encoding axis” that maximally separated movement responses with co-occurring visual signals from those without visual components (Figure 4A, Methods). Activity modes were defined irrespective of retinotopic boundaries – disregarded by movement responses – and averaged across animals using hierarchical bootstrapping (Saravanan et al., 2019) (Methods). In this activity space, single-trial responses across all areas were represented as temporal trajectories, representing the time-varying projections onto the encoding axis (Figures 4B, 4C). Responses with and without stimulus signals were more separated in states of heightened attention (Figures 4B, 4C), and the variability of the projections was reduced. This resulted in overall larger d-prime values when measuring the distributions of response projections in high vs. low states of attention around peak response times (d′ = 3.8 ± 0.7 in high DR trials, d′ = 2.1 ± 0.6 in low DR; mean difference: 1.7, p_boot_ < 0.02, n_boot_ = 50) (Figure 4D, Methods).

**Figure 4.**
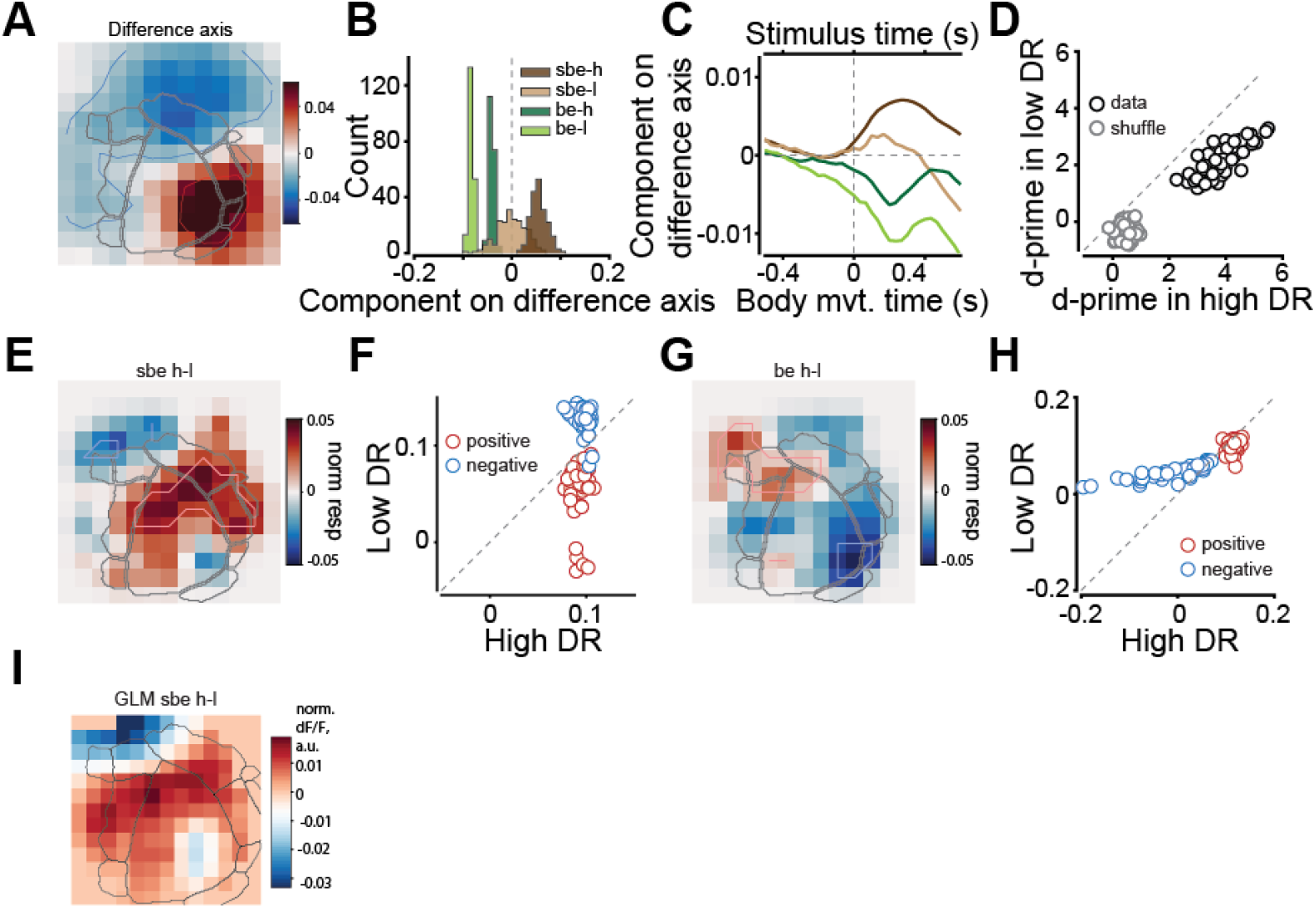
Attention enhances the demixing of visuomotor signals. (A) Activity pattern representing the encoding *axis* that maximally separates movement responses with and without visual components (Methods). (B) Projections onto the encoding axis of saccade-body movement interactions with stimulus components (*sbe*, ‘stimulus + body + eye’ movements) and without (*be,* ‘body + eye’ movements) in high DR (h) and low DR (l) trials across bootstrap resamples for an example mouse (Methods). The d-prime separation of *sbe* from *be* in larger in high DR trials: d′ = 8.06, high DR; d′ = 4.01, low DR. (C) Response trajectories for events as in (B) (example mouse). (D) Population summary for d-prime values in high versus low DR. Each dot is a bootstrap resample. Values are larger in high attentive states (mean difference: 1.7, p_boot_ < 0.02, n_boot_ = 50). Gray dots for the shuffled activity patterns (Methods, mean difference: 0.8, p_boot_ < 0.02, n_boot_ = 50). (E) Normalized difference of activity patterns between high and low DR for the movements-stimulus interactions (Methods). (F) Responses in the positive (red) and negative (blue) regions of the map shown in (E), for the high and low DRs across bootstrap resamples (p_boot_(neg) = 0.04, p_boot_(pos) < 0.02, n_boot_ = 50). (G) Same as (E) for saccade and body movement interactions in the absence of stimulus. (H) Same as (F) for saccade and body movement interactions in the absence of stimulus (p_boot_(neg) = 0.06, p_boot_(pos) = 0.1, n_boot_ = 50). (I) Difference of high and low DR activity patterns as predicted by the DR-dependent partial GLM with only linear stimulus, saccade, and body movement terms. Units are normalized dF/F. Correlation of GLM pattern with pattern directly derived from data in (E) is 0.877 ± 0.024.

To dissect the contribution of region-specific modulations, we computed the difference of the normalized activity patterns for movements with and without stimulus. Normalization removes changes solely due to an overall amplitude modulation, for example, a global multiplicative gain factor, highlighting instead changes in the ‘shape’ of the response patterns. We found that in states of heightened attention, visuomotor activations became larger in regions surrounding the stimulus ROI, significantly in medial V1 and lateral visual areas (p_boot_ < 0.02, hierarchical bootstrapping, n_boot_ = 50), whereas more anterior movement activations were partially suppressed (p_boot_ = 0.04, n = 50, Figures 4E, 4F). Instead, movement activations without visual components showed trends of suppression in V1 and lateral areas (p_boot_ = 0.06, n = 50), with anterior-medial movement regions showing moderately larger activations (p_boot_ = 0.1, n = 50). We also used a shuffle control to test the null hypothesis that these differences emerged spontaneously from amplitude-modulated noise with spatial frequency, as in the data, and found no significant difference between attentional states (Figures S4A, S4B; Methods).

To examine which components were responsible for the pattern change in visuo-motor activations, we analyzed normalized partial predictions of the GLM. Accounting for state modulations of stimulus, saccadic, and body movement responses, and excluding their nonlinear interactions (Methods), closely replicated the change in visuo-motor activity patterns found in the data (Figure 4I) (corr. coef. ρ = 0.88 ± 0.02, mean ± C.I., n = 50). Furthermore, an equally similar pattern change was reconstructed using only stimulus and body movement responses (Figure S4C) (ρ = 0.87 ± 0.02; p = 1.0, two-way ANOVA adjusted for multiple comparisons), whereas pattern changes predicted by all other combinations of components were significantly less similar to the data (p < 10^−4^ for all combinations, two-way ANOVA). Changes in visuo-motor activity patterns with brain state could therefore be largely attributed to state-related changes in visual and body movement responses.

By and large, together with an overall increase in response amplitude, attentional modulations also produced a spatially selective amplification that emphasized the distinctive spatial signatures of visual and movement components.

## Discussion

In this study, we have shown that concurrent sensory, motor, and attentional signals in visual cortical networks interact according to canonical principles of efficient coding: divisive normalization for movement-movement interactions and response demixing for stimulus-movement interactions. Attention enhanced the separation between movement responses in the presence or absence of visual inputs, modulating the underlying neural representations depending on cortical regions.

Divisive normalization in movement-movement interactions can be linked to principles of coding efficiency: the large amplitude of motor signals (about 4 times larger than visual responses) could saturate an information ‘channel’ during signal summation. If the channel is shared with visual inputs, divisive normalization would enable a more efficient use of the dynamical range (Louie and Glimcher, 2019). A contributing factor to divisive normalization might be the mixed response selectivity of individual neurons to saccadic and body movements. If neurons are strongly activated (near saturation) by both saccadic and body movements, then the collective population response to co-occurring events would be larger than for events in isolation, but smaller than their linear sum. The observation that GCaMP responses to individual movements and to their interactions were not saturated does not fully support this interpretation and suggests that other mechanisms might contribute to response normalization (Ferguson and Cardin, 2020).

GLM analysis revealed a supra-linear summation in the interactions between stimulus and movement responses. Although locomotion was absent in our task, this result can be reconciled with studies that examined the effect of locomotion on visual processing at the level of small neuronal ensembles (Dadarlat and Stryker, 2017; Lee et al., 2014; Niell and Stryker, 2010) with response modulations linked to a complex network of cortical and subcortical afferents (Figure S3B) (Lee et al., 2014). In those works, trials with significant locomotion bouts correlated with larger amplitudes of visually-evoked responses. Accordingly, in our widefield data, movement-related activity was significantly detected in the stimulus ROI, summing supra-linearly with visual responses to produce larger visual activations in the presence of movements. However, event-related and GLM analyses also showed that the peak of the activations was not in the stimulus ROI, but rather in the anterior and medial visual areas. Therefore, previously observed response enhancement during locomotion might reflect a strengthening of broadcasted movement signals across all visual areas, but especially along dorsal stream areas, regions that have been implicated in the stabilization of visual perception during movements (Wurtz et al., 2011).

Underlying the demixing of visual and motor components are non-retinotopic principles of visual-area activations followed by movement responses. This signal separation was expected in consideration of the distinct functional and anatomical pathways associated with visual and movement components (Parker et al., 2020). However, this fundamental difference ensures that the demixing is not unique to the set of stimuli used in this study. In all generality, irrespective of the stimuli shown to the animal, the difference between movement modes with and without visual components will always allow defining a non-zero demixing axis. From the viewpoint of the circuits decoding these signals, (linear) separability of visual and motor components observed at these early stages of the visual hierarchy might allow for an efficient use of motor variables in the stabilization of visual perception during movements. Linear separability of these variables has also been reported at the level of small neuronal ensembles in V1 (Stringer et al., 2019) suggesting that demixing of visual and motor signals is a principle that might operate across multiple spatial scales along the visual hierarchy.

Signal demixing was examined here in the context of sustained attention. This form of attention comprehensively relates to arousal, vigilance, motivation, and task-engagement of the animal (Sarter et al., 2001; Unsworth et al., 2018). We adopted this terminology because it is defined in the context of goal-directed behaviors used in this work, although sub-components of sustained attention can also be defined in the absence of a task, for example, arousal. Overall, sustained attention produced a spatially selective amplification of response amplitudes, resulting in a larger demixing of response trajectories in a reduced space of activity modes. The spatial selectivity of the modulation was reminiscent of a center-surround organization, previously observed at the scale of small neuronal ensembles (Zhang et al., 2014). Along the axis that best separated (linearly) movement responses with or without stimulus components, the V1 representation of the stimulus component was stronger around the stimulus ROI and suppressed in ‘surrounding’ movement regions, in anterior-medial areas. GLM analysis revealed that changes in eye movement responses contributed little to this effect, with body movements and stimulus responses contributing the most. Movement patterns without stimulus components changed the least with attentions (besides being overall amplified). Thus, attentional demixing was primarily driven by a *center-surround* change in activity patterns associated with an enhancement of distinctive spatial signatures of visual and body movement responses.

In our task, perceptual and decision-making variables could not be uniquely separated, with movement potentially affecting multiple components depending on their timing within the trial. Because the stimulus was very salient, accumulation of evidence over time should not play a significant role in this task. Decisions could have been made as the percept emerged, right after stimulus presentation, thus with movements affecting both perceptual and decisional components. Later in the trial, it cannot be excluded that mice updated their choice strategy until given the opportunity to make a response, after the end of the open loop. The evidence for this might be related to rotation reversals of the wheel sometimes detected in the open loop and possibly linked to “changes of mind” (procedurally, mice could easily keep the wheel still). Therefore, movements could have affected the decision-making process, also later in the trial. When movements occurred in proximity to the stimulus onset—that is, when they could have affected decisional and perceptual components the most—performance was also the highest. However, fast reaction times were also associated with heightened states of attention. Therefore, a different task design will be necessary to isolate the effect of movements on perceptual and decisional variables, independently from attentional modulations.

In conclusion, our results show that computations linked to the processing of attention-modulated visuomotor signals obey key principles of efficient coding: normalization and activity demixing. These computations allow for the integration of concurrently represented visual and motor variables that remained demixed in a reduced space of activity modes, and with demixing enhanced in heightened attentional states. We speculate that demixing of variables in a low-dimensional coding space should be evaluated in the context of *pattern classification* of neural activity along the convolutional hierarchy of cortical visual areas. In this framework, motor variables can activate visual networks without interfering with perceptual outputs by simply being distinctively classified as non-visual. Furthermore, motor variables can support perceptual stability and reliability by providing these networks with critical contextual information to solve a complex “self-other” disambiguation problem during sensory-motor interactions in complex environments (Cullen, 2004; Feldman, 2009).

## Acknowledgements

We thank Yuki Goya and Rie Nishiyama for their support behavioral training and Rie Nishiyama for her support with animal surgeries. We thank O’Hara and CO., LTD., for their support with the equipment. This work was funded by RIKEN BSI and RIKEN CBS institutional funding, JSPS grants 26290011, 17H06037, C0219129 to AB, and Fujitsu collaborative grant.

## Author contributions

A.B., M.A. and D.L. conceived the study. M.A. collected and pre-processed most of the data, developed the eye tracking toolbox, and analyzed neural and behavioral data. D.L. helped collecting data and did the modeling analyses. R.A. developed the general framework for the behavioral paradigm and helped to collect the data. A.B. supervised all aspects of the work. A.B., M.A. and D.L. wrote the manuscript.

## Methods

### Resource Availability

#### Lead Contact and Materials Availability

This study did not generate new unique reagents. Further information requests should be directed to the Lead Contact, Andrea Benucci.

#### Data and Code availability

Code for analysis and modelling of the data analysis is available at https://github.com/benuccilabncb/cd. Datasets used in this study were deposited to https://github.com/benuccilabncb/cdData.

#### Experimental Model and Subject Details

All surgical and experimental procedures were approved by the Support Unit for Animal Resources Development of RIKEN CBS. Transgenic mice used in this work were Thy1-GCaMP6f mice (n = 10). We appropriately indicated the number of mice when inclusion criteria reduced the number of animals used for specific analysis. For all reported results, the number of sessions per animal ranged from 9 to 60, with a minimum and maximum number of trials per animal from 1000 to 8000. Animals were anesthetized with gas anesthesia (Isoflurane 1.5-2.5%; Pfizer) and injected with an antibiotic (Baytril^®^, 0.5ml, 2%; Bayer Yakuhin), a steroidal anti-inflammatory drug (Dexamethasone; Kyoritsu Seiyaku), an anti-edema agent (Glyceol^®^, 100μl, Chugai Pharmaceutical) to reduce swelling of the brain, and a painkiller (Lepetan^®^, Otsuka Pharmaceutical). The scalp and periosteum were retracted, exposing the skull, then a 4 mm-diameter trephination was made with a micro drill (Meisinger LLC). A 4mm coverslip (120~170μm thickness) was positioned in the center of the craniotomy in direct contact with the brain, topped by a 6 mm diameter coverslip with the same thickness. When needed, Gelfoam^®^ (Pfizer) was applied around the 4mm coverslip to stop any bleeding. The 6mm coverslip was fixed to the bone with cyanoacrylic glue (Aron Alpha^®^, Toagosei). A round metal chamber (6.1mm diameter) combined with a head-post was centered on the craniotomy and cemented to the bone with dental adhesive (Super-Bond C&B^®^, Sun Medical), mixed to a black dye for improved light absorbance during imaging.

### Method Details

#### Behavioral training

Animals were trained in a 2AFC orientation discrimination task. Two oriented Gabor patches (20° static sinusoidal gratings, sf = 0.08 cpd, randomized spatial phase, 2D Gaussian window, sigma = 0.25°) were shown on the left and right side of a screen positioned in front of the animal (LCD monitor, 25 cm distance from the animal, 33.6 cm × 59.8 cm [~58° × 100°dva], 1080 x 1920 pixels, PROLITE B2776HDS-B1, IIYAMA) at ±35° eccentricity relative to the body’s midline. Mice had to report which of the two stimuli matched a target orientation (vertical, n = 6; horizontal, n = 4). The smallest orientation difference varied depending on animals, from 3° to 30°. The largest difference—the easiest discriminations—was ± 90°. Animals signaled their choice by rotating a rubber wheel with their front paws (Figure 1A; Movie S2), which shifted stimuli horizontally on the screen. For a response to be correct, the target stimulus had to be shifted to the center of the screen, upon which the animal was rewarded with 4 μL of water. Incorrect responses were discouraged with a prolonged (10 s) inter-trial interval and a flickering checkerboard stimulus (2 Hz). If no response was made within 10 s (time-out trials), neither reward nor discouragement was given.

Animals were imaged after exceeding a performance threshold of 75% correct rate for 5–10 consecutive sessions (typically after ~4–12 weeks) when trained in the automated self-head-restraining setups. Depending on animals, performance in the imaging setup (e.g., Figure 3E) could fluctuate from session to session. To work with a coherent behavioral dataset, we excluded sessions with exceedingly large fractions of time-outs (>= 20%) or with average performance dropping below 60%.

Every trial consisted of an open-loop period (OL: 1.5 s) and a closed-loop period (CL: 0–10 s), followed by an inter-trial interval (ITI: 3–5 s randomized). We recorded cortical responses, wheel rotations, and eye/pupil videos from a pre-stimulus period (1 s duration) until the end of the trial. Stimuli were presented in the OL period, when wheel rotations did not produce any stimulus movement. In 25% of the trials, the OL lasted longer by an additional randomized 0.5–1.5 s period, during which we presented simulated-saccade stimuli (i.e., grating stimuli moving passively on the screen according to the previously recorded eye movement velocities) (Figure S1G).

#### The psychometric curve

We fitted the animal’s probability of making a right-side choice as a function of task difficulty using a psychometric function *ψ*(*ϵ*; *α*, *β*, *γ*, *λ*) = *γ* + (1 − *γ* − *λ*) *F*(*ϵ*; *α*, *β*), where *F*(*x*) is a Gaussian cumulative probability function, α and β are the mean and standard deviation, *γ* and *λ* are left and right (L/R) lapse rates, and *ϵ* is the signed trial difficulty. Confidence intervals were computed by bootstrapping (n = 999).

#### Detection of saccades and body movements

##### Eye tracking

We monitored the left contralateral eye illuminated by IR LED (SLS-0208-B medium Beam, Mightex^®^), using a CMOS camera (FL3-U3-13E4M-C, POINT GREY) equipped with a zoom lens (Navitar Zoom 7000, 1280×1024 pixels, typical ROI size: 350×250 pixels, 30Hz acquisition rate) with an IR filter (Kenko PRO1D R72, 52mm). The camera was aligned to the perpendicular bisector of the eye, making ~60° angle with the midsagittal axis of the animal.

Automatic tracking of the pupil position was done with custom software (Matlab, Mathworks^®^). We first processed each video frame to extract the visible region of the eye ball (MATLAB *imreconstruct* and factorization-based texture segmentation), with morphological operations (dilation, erosion, disk structuring elements 106 and 202 μm, respectively) to remove pixel noise. To extract the pupil segment, which has lower intensity values, we performed Otsu thresholding on the intensity distribution in every frame. We further imposed geometrical constraints to reduce misclassification of the pupil with the eyelid shadows: the pupil had to 1) be closer to the center of eye segment (Euclidean distance) and 2) have a roundness index (4*pi*area/perimeter^2^) >0.7. We fitted an ellipse to extract the pupil center position and area, and then used it for saccade detection and pupil area analyses. We also confirmed the accuracy of pupil tracking by visually inspecting hundreds of trials.

##### Saccade detection

To detect saccadic eye movements, we first filtered the XY positions of the pupil center over time (frames) using an edge filter [−1 −1 0 1 1] and transformed the resulting time series to XY velocities. We then applied an adaptive elliptic thresholding algorithm to find the saccade time frames that had velocities larger than the elliptic threshold. We discarded the saccades that lasted <= 60 ms and were smaller than 1.5° (see ERA method-section for the robustness of the results relative to specific threshold values). We extracted the time, magnitude, duration, velocity, start, and landing positions of each saccade (Figure 1C).

##### Pupil Area

To analyze the pupil area (Figure 3C), we first converted eye-tracking-camera pixels to mm using direct measurements of the width and length of the eye to account for experiment-to-experiment variability in the zooming factor. We calculated the average pupil area for each imaging session by averaging area values across all trials within the session. Lastly, pupil area in every trial was normalized (subtracted) relative to the session mean.

##### Wheel detection

We automatically detected the time at which the animals rotated the wheel. We flagged as potential wheel movement time points when the velocity had a zero-crossing (i.e., sign change) and deviated from zero above a fixed threshold (20°). Movements smaller than such threshold were considered unintentional twitches of the wheel and discarded (see ERA method section for the robustness of the results relative to specific threshold values).

#### Imaging

Expert mice were placed under a macroscope for wide-field imaging (THT, Brain Vision) using a head-plate latching system. A tandem-lens epifluorescence macroscope was equipped with a CMOS camera (pco.edge5.5, pixel size: 6.5 μm^2^, pixel number: 5.5 mp) and two lenses (NIKKOR, 50 mm, F1.2, NA = 0.46) to image GCaMP6f fluorescent signals: excitation light, 465 nm LED (LEX2B, Brain Vision); emission filter, band-pass at 525±25 nm (Edmund).

##### Retinotopy

We computed maps of retinotopy to identify primary and higher visual areas. Briefly, we used a standard frequency-based method with slowly moving horizontal and vertical flickering bars in anesthetized mice (~0.8% Isoflurane) on a 40” LCD monitor (Iiyama^®^). Visual area segmentation (Figures 1D, S1C) was done based on azimuth and elevation gradient inversions. To center and orient maps across animals, we used the centroid of area V1 and the iso-azimuth line passing through it.

##### Pre-processing wide-field GCaMP6f signals

We first motion corrected GCaMP data. Using a semi-automated control-point selection method (MATLAB *cpselect*, using blood vessel images), all image frames were registered to a previously acquired retinotopic map. To compute relative fluorescence responses, we calculated a grand-average scalar 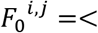 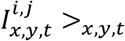, with 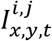 the XYT image tensor in trial *i*, session *j*. We then used this scalar to normalize the raw data tensor 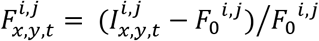. The data in each trial were then band-pass filtered ([0.1 12] Hz) and smoothed with mild spatial filtering (Gaussian σ = 20 μm). Lastly, each tensor was compressed with spatial binning (130×130 μm^2^ with 50% overlap). The results presented do not critically depend on any of these parameters.

#### Imaging Data Analysis

##### Event-related analyses (ERA)

We analyzed isolated events in windows that contained only one of the four events: stimulus, simulated saccade, saccade, and body movements (Figures 1E, S1D, S1E, S1G, S1H). The stimulus isolation window was from 1 s before to 1 s after the stimulus onset; the simulated saccades window was from trial start to 3 s after stimulus onset; and the saccades and body movement window was from 2.5 s before to 2.5 s after the event. The window sizes were chosen by considering the time needed for the response to return to baseline during a quiescence period. We also excluded trials when other events were detected 2.5 s away from the closest event on each side.

For event-related analysis (ERA), we computed trial-averaged responses centered on the time of the event. Spatially, we defined four ROIs for each event: we first identified the time of peak response amplitude in V1 and then selected pixels above a varying threshold, from 70^th^ to 99^th^ percentile at steps of 0.5 percentiles, to create binary mask images. We then averaged the masks and defined an ROI as a contiguous group of pixels above the 99^th^ percentile. The results presented did not critically depend on any of the parameters above. Temporal event-related responses in each ROI were computed as a within ROI pixel average after frame-0 correction. This was done by computing an average dF/F in a time window [−0.2, 0] s from stimulus onset and simulated saccade, or [−0.8, −0.3] s from saccade and body movements, averaged across trials and animals, and subtracting this value from the event-related responses. Error bars in across-animal averages are standard error of the mean (s.e.), and across-trial error bars are at a 95% confidence interval (95% CI). Peak responses were computed by averaging within a 100 ms window centered at the time of max amplitude. To compute spatial maps (Figure 1E, and maps in Supplementary Figures), we normalized (z-scored) the dF/F of each pixel in every frame with max amplitude over time: 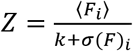, where *F*_*i*_ is the peak amplitude (average of peak frame ±1 frame) on trial *i*, 〈*F*_*i*_〉 is the average across trials, *σ* is the standard deviation across trials, and *k* is a small regularizing scalar to avoid division by zero. Then, we averaged the z-scored responses across mice (Figure 1E).

##### Linear prediction with jittered times

To compute the linear prediction for stimulus-movement interactions (Figure 3A), we convolved the isolated movement responses (Figure 1E) with binary input vectors representing recorded movement times, summed them with the isolated stimulus response, and averaged across trials and animals. Similarly, for saccade-body movement interactions (Figure 3A), we convolved the responses to isolated body movements and saccades with the corresponding binary input vectors, aligned them to the time of saccade, and averaged across trials.

##### Signal Saturation

For a given trial *i* with a pair of saccade and body movement events having a time lag in the range [−0.25, 0.25] s, we calculated baseline fluorescence in primary visual cortex (V1), 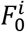, by averaging raw fluorescence values over t = [−0.8, 0] s from stimulus onset and averaging across all V1 camera pixels. Similarly, we calculated the peak fluorescence for the V1 area, *F*^*i*^, by averaging raw fluorescence values over a 100 ms window centered at the time of the peak interaction response and averaging across all V1 camera pixels. The percentage amplitude change was defined as 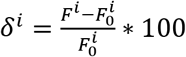. Then we divided the distribution of 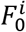 derived from all trials into five equal-amplitude intervals (quintiles), and for each interval, computed mean 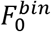 values together with corresponding mean percentage changes *δ*^*bin*^. For every animal, we plotted *δ*^*bin*^ as a function of 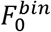 with its 95% CI (gray lines in Figure S2A, middle). We discarded intervals with less than 25 trials. To average across animals, we further binned 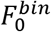 and calculated the mean and s.e. for each bin (black line in Figure S2A, middle).

##### DR, performance, pupil area, and reaction times

To define a *pupil space* (Figure 3C), we calculated for each trial the baseline pupil area, x-axis (i.e., the average area in the [0, 200] ms interval after the stimulus onset relative to the session average), and on the y-axis, the maximum area change relative to this baseline in the OL period (Figure 3C). For interacting saccadic-body movement events, the baseline was computed as the pupil area at the time of the first detected event (±50 ms), and the maximum change was calculated in a [0, 4] s interval after the second event. The dynamical range index (DR) was calculated using the standard deviation of the V1 response (average across all V1 pixels) over the whole trial duration *DR*_*i*_ = *σ*[*R*_*i*_ (*t*)] where *R* is the average V1 response in trial *i* over time *t*. Calculating DR values, including responses from other areas, did not significantly change these results. To define ‘high’ and ‘low’ DR states, we used 70th and 30th percentiles of DR distribution across all trials. Reaction times were defined as the average detection times for saccade and body movements after the stimulus presentation (Figures 3B, 3D).

#### Cortical state: correlations

To test for the generality of the attentional effects relative to the definition of the DR index, we computed a second measure of cortical state based on response correlations (Figure S3A). In every trial, we calculated the variance of each image pixel, 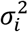, and the covariance across all pixels, *Coν*_*i,j*_. The cortical state (C) for a given trial was calculated as the overall variance in the population 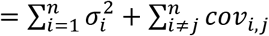, with *n* the total number of pixels. Trials were labeled as being in a high, low, or middle cortical state based on their *C* values relative to the overall C-distribution (> 70^th^ percentile, < 30^th^ percentile or > 30^th^ and < 70^th^ percentiles, respectively).

#### Generalized linear model

##### Data used in GLM

In each trial, for every pixel, GCaMP responses were frame-zero corrected by subtracting the average dF/F in the [−1.0, −0.8] s interval before stimulus onset. Data was downsampled to 10 Hz and spatially binned: 300 x 300 μm pixel size, with a bin referred to as “tile” in the following. Only responses in the open loop (non-interactive period) were analyzed to exclude activations due to stimulus motion. Trials with events in the [−1.0, −0.8] s interval before stimulus onset or with blinks or simulated saccades were excluded. We did not have sufficient statistical power to examine higher order interactions, that is, interactions between multiple wheel and eye events, whose effects are included in the 2nd order wheel-eye kernels.

##### Model design

For a given tile and trial, we model the GCaMP response *y*(*t*) as 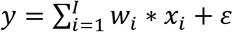, with convolutional kernels *W*_*i*_, Gaussian noise *ε*~*N*(0, Σ), and inputs *x*_*i*_, *i* ∈ {*s*, *b*, *e*, *sb*, *se*, *be*}, with *s*, *b*, *e*, stimulus onset, body movement, eye movements, and their pairwise combinations, *sb*, *se*, *be*. Each *x*_*i*_ was a binary time series, with 1’s at the time of an event. Pairwise inputs were the outer product of corresponding linear inputs. Kernels *W*_***i***_ acted causally and anticausally to account for both pre- and post-movement responses. The bias term was zero since *y* was frame-zero corrected.

##### Optimization

In matrix form *Y* = *XW* + *εI*; we estimated kernels from 40 data bootstraps using ridge regression, 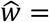 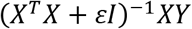, where the optimal *ε* is found for every tile and kernel *W*_*σ*_ by maximizing log marginal likelihood using a fixed-point algorithm^73,74^. The expression for 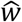 is equivalent to the Bayesian MAP estimate with 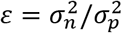, where 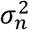 is noise variance of observations and 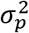 is prior variance^75^ 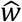 biased, with the amplitude of kernels estimated from relatively few noisy trials strongly penalized.

##### Sequential fitting

To eliminate the trade-off between kernels of different inputs, we estimated them sequentially. We estimated *W*_*s*_ from trials with no body or eye movements until 2.8 s after trial start, *W*_*s*_ was estimated in a time window *τ*_*s*_ = (−1.0, 1.5) s centered on the stimulus onset and could also contain a slow upward/downward trend related to movements in the ITI period. From the residuals, *y*_*rs*_ = *y* − *W*_*s*_ ∗ *x*_*s*_, we estimated *W*_*e*_, with *τ*_*e*_ = (−0.3, 2.0) s and *W*_*b*_ with *τ*_*b*_ = (−0.3, 2.0) s using segments of trials where the eye and body movements were isolated. Isolation meant no overlap with any part of the *τ*-window of any surrounding movements. From the residuals, *y*_*rsbe*_ = *y*_*rs*_ − *W*_*b*_ ∗ *x*_*b*_ − *W*_*e*_ ∗ *x*_*e*_, we estimated the body-eye movement interaction kernel *W*_*be*_, *τ*_*be*_ = [(−0.3, 2.0) s, (−0.3, 2.0) s] using all trials. Lastly, we estimated stimulus-eye movement *W*_*se*_ (using dF/F downsampled at 5 Hz) and stimulus-body movement *W*_*sb*_ kernels from the residuals *y*_*rse*_ = *y*_*rs*_ − *W*_*e*_ ∗ *x*_*e*_ and *y*_*rsb*_ = *y*_*rs*_ − *W*_*b*_ ∗ *x*_*b*_, using the same trial segments as when fitting *W*_*e*_ and *W*_*b*_ respectively to ensure isolation. Baseline of *y*_*rsb*_ of every tile was additionally aligned to its mean in (−0.5, −0.3) s before the stimulus onset to further minimize the influence of ITI movements, and only trials with body movements at the time or after stimulus onset were used to estimate *W*_*sb*_ to avoid baseline correction to pre-motor activity.

##### Explained variance

We estimated the response variance of every tile of every animal explained by a full GLM using 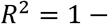 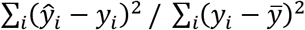, where 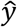 is GLM prediction and 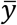 is data average, with summation done over individual time bins and trials, following a 5-fold cross-validation procedure. We report population average maps of explained variance in percent units (Figure S2B).

#### Kernel analysis

##### Used data

Despite the stringent trial-selection criteria, *W*_*be*_ could be reliably estimated from n = 8 animals, *W*_*se*_ from n = 5 animals, and *W*_*sb*_ from n = 6 animals. *W*_*se*_ and *W*_*sb*_ could be estimated in fewer animals than *W*_*be*_ because we additionally required isolation of the respective movement.

##### Graphical representation and pre-processing

We represent kernels *W*_*be*_, *W*_*se*_, *W*_*sb*_ in the coordinates of lags, (*τ*_*b*_, *τ*_*e*_), (*τ*_*s*_, *τ*_*e*_), (*τ*_*s*_, *τ*_*b*_) (Figures 2D–2F), a kernel element is thus, for example, *W*_*be*_ (*τ*_*b*_, *τ*_*e*_). For improved graphics, we filtered *W*_*be*_, *W*_*se*_, *W*_*sb*_ in the lag-lag space with a mean filter of 3×3 time bins. For the results presented, we only considered elements significantly different from zero (two-tailed Mann-Whitney U-test at *α* = 0.05), that passed the shuffle test (*W*_*be*_ only), and that could be estimated from at least 10 data points—these criteria were tested for all animals, all tiles, and lags. The shuffle test for *W*_*be*_ was performed by randomly assigning trials with interactions to saccade-body movement pairs with different Δ*τ* and fitting the GLM with *ε* fixed at the unshuffled estimate. *W*_*be*_ was then tested against the shuffled-data estimate of *W*_*be*_ using Mann-Whitney U-test (*α* = 0.05).

##### Population kernels

We calculated population kernels 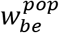, 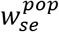, or 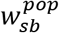 (Figures 2D–2F), as average normalized kernels *W*_*be*_, *W*_*se*_, *W*_*sb*_ belonging to the tiles (300×300 μm) with the largest negative - *W*_*be*_(*τ*_*b*_, *τ*_*e*_) – or largest positive - *W*_*se*_(*τ*_s_, *τ*_*e*_) and *W*_*sb*_ (*τ*_*s*_, *τ*_*b*_) - values. The patterns of nonlinear contribution of *W*_*be*_, *W*_*se*_, *W*_*sb*_ did not change substantially as we considered different larger regions (data not shown). We masked elements 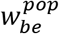 (dim colors) that were indistinguishable from permuted data or could not be estimated in n = 3 or more (of 8 total) animals. We masked elements 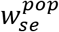 and 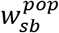 if they could not be estimated in n = 2 or more (of 6) animals.

We show (*τ*_*b*_, *τ*_*e*_) of maximally negative elements of every animal with a red dot, and population average 〈(*τ*_*b*_, *τ*_*e*_)〉 with a red circle (Figure 2E), and s.e. smaller than circle size. We excluded one outlier mouse. Similarly, we show the largest positive *W*_*se*_ (*τ*_*s*_, *τ*_*e*_) of individual animals with black asterisks and median ± median-derived s.d. as a circle with error bars (Figure 2D). We show largest positive *W*_*sb*_ (*τ*_*s*_, *τ*_*b*_) of individual animals as green crosses, and a population median and median-based standard deviation as a large cross (Figure 2D). Markers of all animals were jittered by < 1 kernel bin width to reduce overlap in the graphics. Median-derived s.d. was computed as 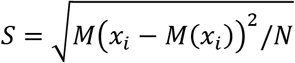 where *M* is median operator, *N* – number of animals, *i* – animal index, and *x*_*σ*_ stands for *τ*_*s*_, *τ*_*b*_, or *τ*_*e*_ of maximum of *W*_*sb*_ or *W*_*se*_ of animal *i*.

##### GLM predicted responses

We predicted nonlinear components of the response using the GLM (Figures 2D–2F), where all but the corresponding nonlinear term was set to zero. Responses were generated according to lags highlighted on the respective population kernels.

#### Normalization model

This analysis was based directly on the data averages (*W*_*de*_, *W*_*db*_, *W*_*ds*_) marked with a subscript “d” to distinguish them from GLM kernels (*W*_*e*_, *W*_*b*_, *W*_*s*_); these data averages were convolved with respective inputs to obtain response components (*r*_*de*_, *r*_*db*_, *r*_*ds*_). We used the same mice as were used for GLM analysis (n = 8 out of 10).

We considered a normalization model of co-occurring saccade and body movement responses in which the magnitude of their normalization at a given time was proportional to the product of their magnitudes. Total model response 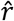 was thus equal to 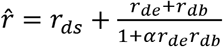, where *α* is the normalization parameter, which we find as 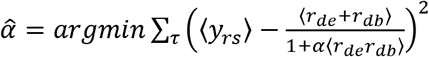, where *y*_*rs*_ = *y* − *x*_*s*_ * *W*_*s*_ is a residual after subtracting the stimulus component of the GLM from the recorded response, and where *τ* ∈ [0, 0.5] s relative to the saccade, 〈·〉 is averaging across trials, and fitting is done with (n = 20) bootstrap resamples of data, using MATLAB *fmincon* in the range [−5, 20]. We made a prediction of interacting saccade and body movement responses in all trials from the fitted normalization model, 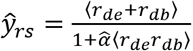, and used these predictions to fit the interaction kernel *W*_*be*_ of the GLM (Figure 2F), given that by design 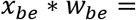 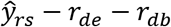. We used this *W*_*be*_ to generate spatial activity profile at the time of maximum negative nonlinearity (Figure 2F, right).

#### Activity modes, encoding axis

Wide-field data: We aligned imaged cortices from all mice using retinotopic boundaries and for each mouse selected trials with movements and no stimulus components according to the following criteria: eye movements had to occur within ±0.75 s from wheel events. The 1^st^ movement had to be at least 1.5 s away from stimulus onset, and the 2^nd^ event separated by at least 0.25 s from any other future movement. Movements with stimulus components had to occur within 1.5 s from stimulus onset, but not before 0.2 s after stimulus onset, allowing for the stimulus response to become detectable before movement onset. Movement responses without stimulus components were frame-zero corrected, averaging frames in a 0.3 s window centered ~1 s before the wheel detection time, and the average activation patterns were computed by averaging frames in a [0, ~0.25] s window. Movement responses with stimulus components were frame-zero corrected averaging frames in a 0.2 s window centered ~1 s before the stimulus detection time, and activation patterns were computed by averaging frames in the [~0.2, ~0.35] s window after stimulus. Average activity patterns were normalized (vector norm), and their difference defined the encoding axis shown in Figure 4A. Unresponsive tiles (with response standard in the lower 20^th^ percentile of the distribution across tiles) were excluded, typically near the edges of the implanted chamber. This procedure was first applied to all trials, irrespective of attentional state (all DR conditions). Selecting only medium DR trials for this step did not significantly affect the results. We then applied the same procedure separately to high or low DR trials using hierarchical bootstrap across animals and trials (see Methods below). Each average activity pattern was normalized using the normalization factor applied in computing the encoding axis. Projections onto the encoding axis were the dot-product between vectors. Thus, changes in projection amplitudes onto the encoding axis represented modulations relative to a grand average across all DR states. Instead, to examine DR-dependent changes in response patterns, aside from overall amplitude modulations (e.g., multiplicative gain scalar), Figures 4E–4H), we normalized movement with or without stimulus components in different DR states using their own vector norm. This procedure highlighted changes in spatial ‘structure’, disregarding global amplitude modulations.

To examine which activity components were responsible for the DR-dependent response pattern change (Figures 4I, S4C), we made partial predictions from the GLM. Here, GLM was separately fitted on high and low DR trials, and DR-dependent responses on individual trials were predicted using only the stimulus, saccade, and body movement components without their interactions, and using stimulus and body movement components. Predictions from other components were also made (all possible individual and pairs of components), but their correlation with the data pattern was significantly lower (data not shown). We followed the same hierarchical bootstrapping procedure for partial GLM predictions as for the above analyses.

The d-prime projection values onto the encoding axis generated four distributions: two distributions for movement with or without stimulus components and two distributions for the high and low DR states. We calculated d-prime as: 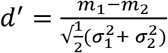 with *m* and *σ* mean and standard deviation of the corresponding pair of compared distributions.

#### Shuffle control

The shuffle control in Figure S4 was implemented within the hierarchical bootstrap loop. Right before the projection step, the (i,j) indices of the camera frames were randomly shuffled, and then a Gaussian spatial filter was applied to match the characteristic spatial frequency of the unshuffled data (*σ* = 1/10 of the frame size). Lastly, the dynamical range of the frame ([min, max]) was also matched to that of the data.

#### Hierarchical bootstrap

We applied hierarchical bootstrap (HB) to merge data across animals. HB is used here to account for possible ‘nested’ hierarchical correlations within datasets from individual animals (Saravanan et al., BioRxiv 2019). To this end, we first created a bootstrap loop for the animals, and at each iteration, we also bootstrapped trials within each animal. We used n = 50 bootstraps for animals and m = 100 bootstraps for trials. Doubling these numbers did not significantly affect the results. At the end of each bootstrap iteration, we computed the relevant statistics, d-prime values, for high and low DR states. Bootstrap confidence intervals were the 5^th^ and 95^th^ percentile of the bootstrapped distribution. We also report p-boot (*p*_*boot*_) values computed as the proportion of means in the population of bootstrapped means that were greater or equal in high DR than low DR states, setting a false positive rate α = 0.05.

### Quantification and Statistical Analysis

Details of the analyses can be found in the corresponding sections of the results, in figure legends, and STAR Methods. We used confidence intervals of the mean (C.I.) for within animal confidence statistics. We used standard error of the mean (s.e.) for across animals error estimates. We used a t-test to compare mean amplitudes from within-animal data. When pooling maps across animals, we first z-scored and then averaged. We used the term ‘Wilcoxon’ to refer to the Wilcoxon signed-rank test, and ‘U-test’ to refer to the Wilcoxon rank-sum test. Significance was established at a p-value of 0.05, adjusted for multiple comparisons using Bonferroni correction in cases where it was necessary.

