## Supplemental information for "Attention Decorrelates Sensory and Motor Signals in the Mouse Visual Cortex"

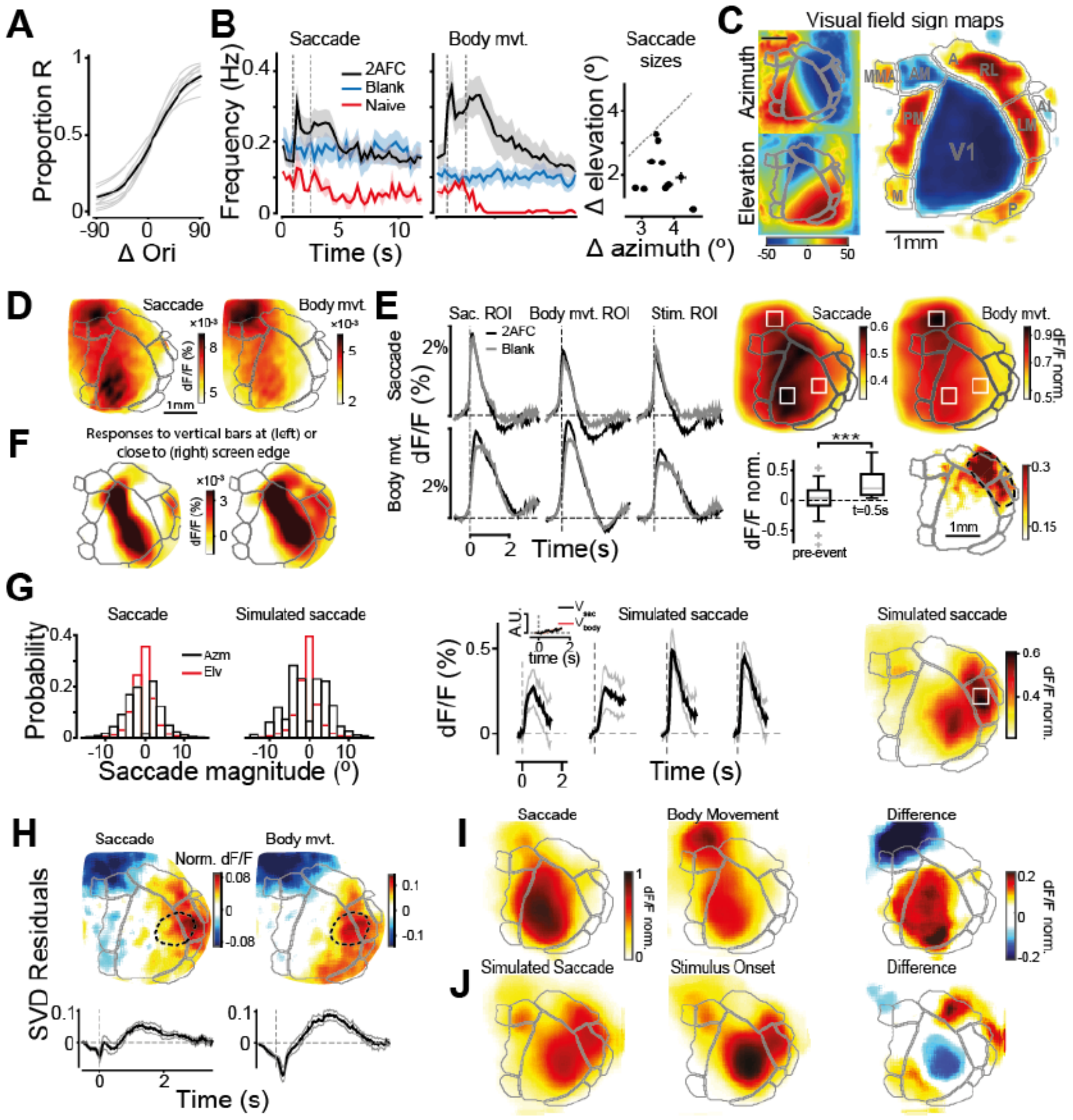


**Figure S1**. **Distinctive behavioral and neural patterns during the task. (A)** Psychometric curves. Black line: average across animals (n = 9). Sessions included had average performance > 60 % and time-out rate < 20 % (n = 9); only one mouse did not reach 60 % performance. Light gray lines for individual mice. Abscissa, task difficulty: ± 90° for the easiest conditions when either the left or right stimulus is the target (vertical, n = 6 mice; horizontal, n = 4 mice). **(B)** Frequency of saccades (left) and body movement across trial time (middle) in 2AFC (black) and 'blank' experiments (blue, passively head-fixed mice in front of blank screen). Red line, naive mice starting to be trained in the same 2AFC tasks (n = 2). Vertical dotted lines, open-loop period; shaded bands, s.e. Right. Average azimuth and elevation changes of saccades, dots are averages across all detected saccades for each animal. **(C)** Average azimuth, elevation, and sign maps aligned across mice. Gray lines, segmented visual area borders. **(D)** Pre-event activity pattern for saccades (left, t = [-100, 0] ms) and body movements (right, t = [-100, 0] ms). **(E)** Body movement and saccade related responses during blank condition (no visual stimulus and no task). Left: Responses to saccades (top) and body movements (bottom) in 3 different ROIs (columns; shown on the rightmost maps), averaged across mice examined in both blank and 2AFC conditions (n = 5; peak response in blank versus 2AFC, t-test, p > 0.5.

FWHM was also unchanged in 2AFC vs blank conditions, t-test, $p > 0.5$). Right-top: map of saccadic and body movement responses during blank experiments. Compare to Figure 1E. Right-bottom: difference between 2AFC vs blank saccadic response amplitudes before ([-0.27 -0.13] s) and after (at t = 0.5 s) saccade detection ($n = 6$, $p < 10^{-3}$) with a spatial localization in motion sensitive areas (dashed contour line). **(F)** Responses to a vertical flickering bar. Spatial frequency: 0.07 cpd; temporal frequency: 0.5 Hz; Size: 15° of azimuth and 80° of elevation, with azimuth corresponding to the contra-lateral vertical edge of the screen; left panel: 45° relative to the body midline; right panel: 30° relative to the body midline. **(G)** Left: Probability distributions for saccade azimuths and elevations (left histograms), randomly sampled to generate simulated saccades (right histograms, Methods). Middle: Response to simulated saccades in different ROIs (from left to right: saccade, body movement, simulated saccade and stimulus ROIs). The ROI for simulated saccade is shown as a white square on the rightmost map. Peak simulated saccade response: 0.53 ± 0.04 %. Right-most map: peak of the simulated saccade response. **(H)** Map of residuals from an SVD decomposition of movement responses (Methods) estimated using the time point of peak residual amplitude in the stimulus motion ROI (dotted contours; compare to the rightmost map in panel g; t = 1.4 s and t = 1.7 s for saccade and body movement). Time series of the residuals in the stimulus motion ROI are shown below the maps. Maximum normalized residual amplitude for saccades: 0.05±0.01 (s.e., t-test $p < 0.01$; $n = 10$) and for body movements: 0.09 ± 0.01 (s.e., t-test $p < 0.001$). All dF/F values reported here are normalized to the peak response (Methods). **(I)** Average response patterns at the time of peak amplitude for saccades and body movements. **(J)** Same as (I) for simulated saccades (panel (G)) and for stimulus onset.

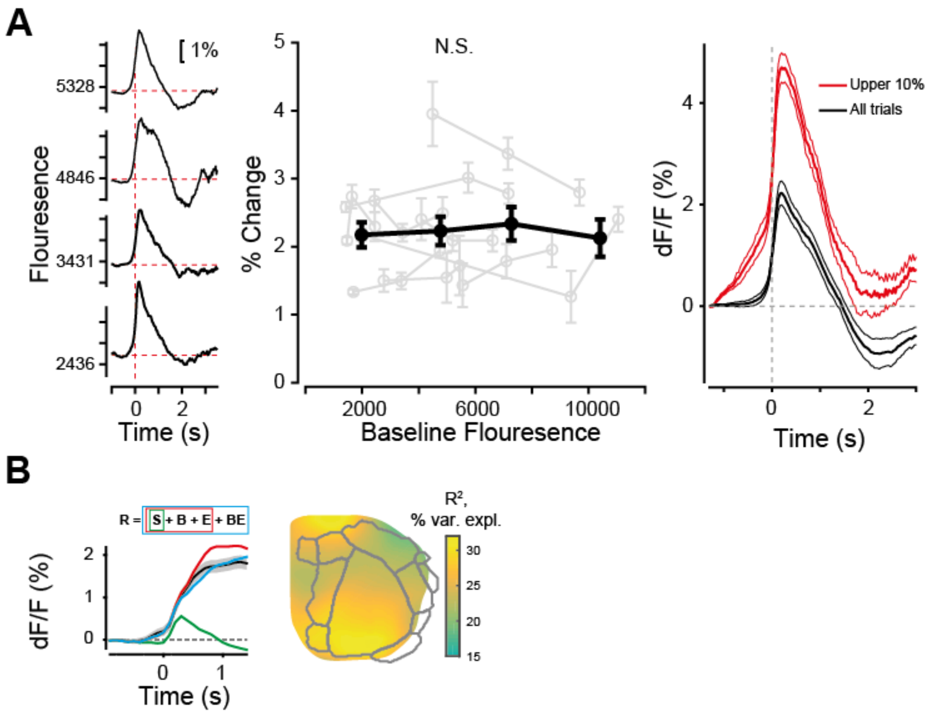


**Figure S2. Signal saturation analysis and response amplitude reduction during co-occurring saccades and body movements (model). (A)** Left: example interaction responses showing the % change at different levels of baseline fluorescence (rows). Middle: Peak % change as a function of baseline fluorescence, for each mouse (grey) and their average (black). Right: peak response values can significantly exceed amplitude values shown in Figure 1E (red line, trials in the upper 10% of the distribution of peak response amplitudes; black line, average across all trials). **(B)** Left. GLM sequential fitting: average dF/F aligned to the stimulus onset in trials with saccade-body movement interactions (black), gray band C.I., example mouse. Model responses: stimulus (green), stimulus + isolated movements (red), stimulus + isolated movements + nonlinear saccade-body movement interaction (blue). Right. Average variance explained ($R^2$) by the full GLM model with interactions (n = 8).

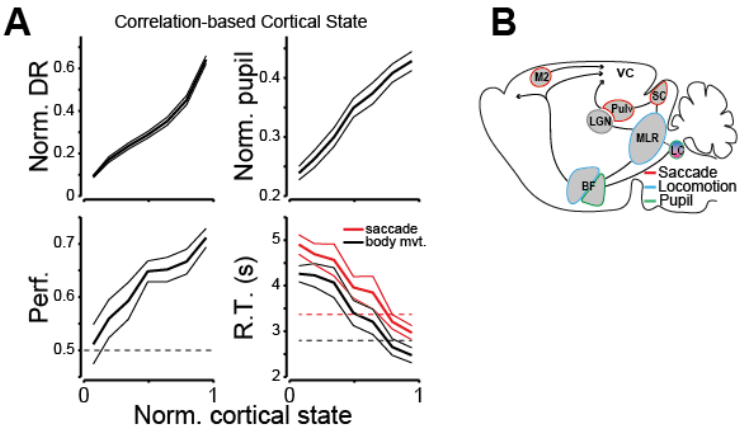


**Figure S3. Correlates of high and low DR index trials. (A)** Cortical states defined from response correlations (Methods) are correlated with DR index ($p < 10^{-5}$, t-test), pupil area ($p < 10^{-5}$), performance ($p = 4.8 \cdot 10^{-5}$) and reaction times ($p < 10^{-5}$, both saccade and body movement R.T.). **(B)** Schematic of known circuits involved in the control of saccades, body movements, and pupil-related arousal states. Their differential recruitment might mechanistically explain response modulations: VC, visual cortex. Pulv, pulvinar. SC, superior colliculus. LGN, lateral geniculate nucleus. MLR, mesencephalic locomotor region. LC, locus coeruleus. BF, Basal forebrain. M2, secondary motor cortex.

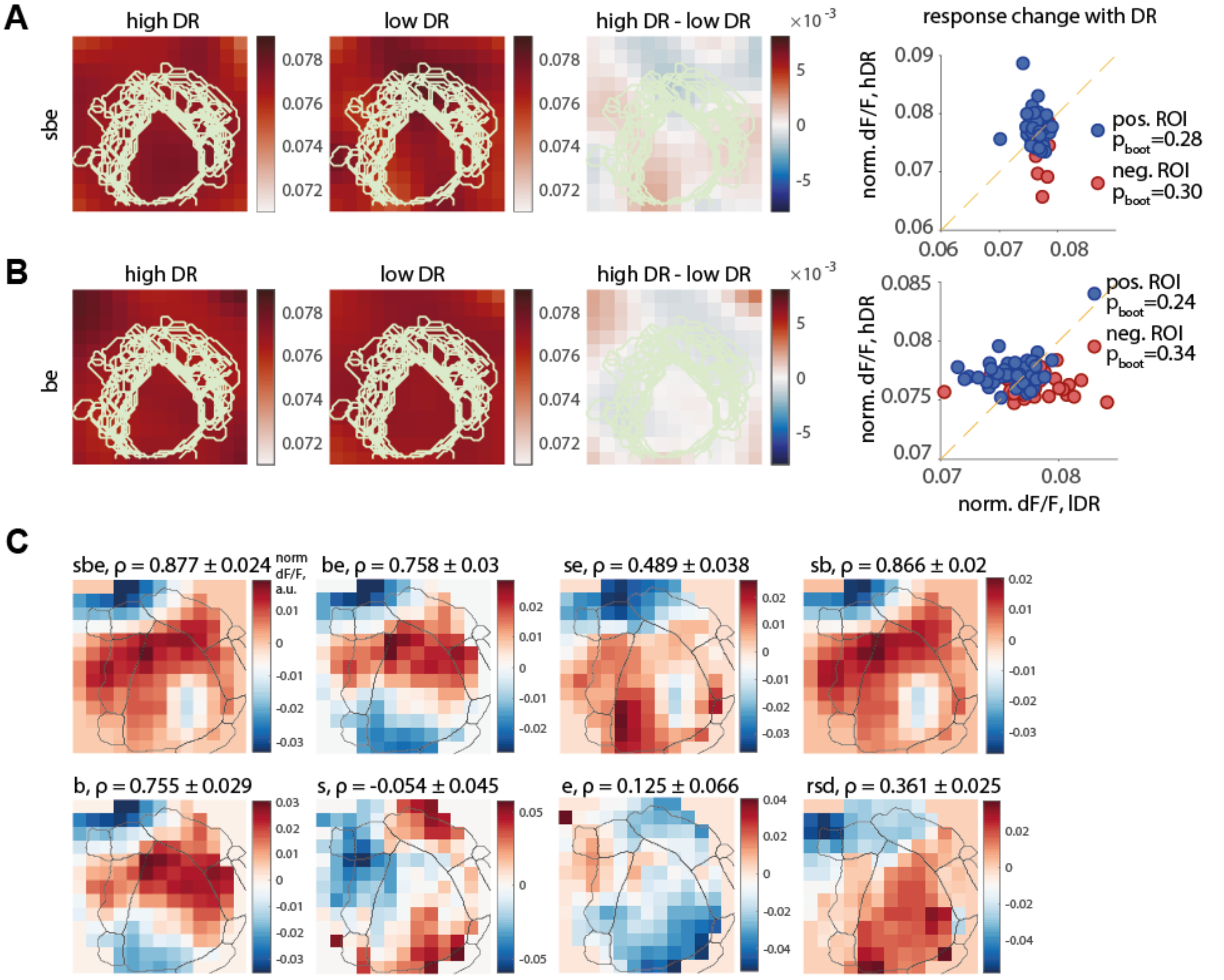


**Figure S4. Shuffle controls for d' analysis and pattern changes obtained from GLM. (A)** Activity patterns for movements with stimulus signals (stimulus + body + eye movements – sbe) are computed separately in high and low DR trials (left and middle panel) together with their difference (right panel). Data was shuffled at the pixel level, keeping the characteristic spatial frequency of the unshuffled data and its dynamical range (Methods). This procedure controls whether the difference patterns observed in Figure 4 emerge from averaging noisy responses with characteristic spatial frequency as observed in the data. Rightmost panel, significance of the most positive and negative regions tested (same procedure used in Figure 4). The color of the dots reflects the response sign as shown in the colorbar; no region reached significance. **(B)** same as in (A), but for body + eye movements – be. **(C)** Difference of the normalized activity patterns as predicted by the GLMs that included only a subset of activity components (in titles): *sbe* - stimulus, body movement, eye movement, *be* – body and eye movements, *se* – stimulus and eye movements, *sb* – stimulus and body movements, *b* – body movements, *s* – stimulus, *e* – eye movements, *rsd* – residual of the recorded activity on every trial and *'sbe'* for predictions. All components of the GLM were computed in the high and low DR trials separately and were used to predict activity on trials with corresponding DR values. Hierarchical bootstrap procedure is used as in Figure 4. ρ values in the titles – Pearson correlation of GLM pattern differences with data pattern difference (Figure 4E), C.I. over n = 60 bootstraps. Difference between the ρ(*sbe*) and ρ(*sb*) not significant, 2-way ANOVA corrected for multiple comparisons, all other correlations are significantly smaller, $p < 10^{-4}$.
